# SIRT1 ameliorates premature senescence-induced defenestration in hepatic sinusoidal endothelial cell

**DOI:** 10.1101/2020.04.24.059048

**Authors:** Xiaoying Luo, Yangqiu Bai, Shuli He, Xiaoke Jiang, Zhiyu Yang, Suofeng Sun, Di Lu, Peiru Wei, Yuan Liang, Cong Peng, Yaru Wang, Ruli Sheng, Shuangyin Han, Xiuling Li, Bingyong Zhang

## Abstract

Premature senescence, linked to progerin, involves in endothelial dysfunction and liver diseases. Activating sirtuin 1 (SIRT1) ameliorates liver fibrosis. However, the potential mechanisms of premature senescence in defenestration in hepatic sinusoidal endothelial cells (HSECs) and how SIRT1 affects fenestrae remains elusive. Our study showed that *in vivo*, premature senescence occurred, with decrease of SIRT1, during CCl_4_-induced defenestration in HSECs and liver fibrogenesis; whereas overexpressing SIRT1 with adenovirus vector lessened progerin-associated premature senescence to relieve CCl_4_-induced defenestration and liver fibrosis. *In vitro*, fenestrae in HSECs disappeared, with progerin-associated premature senescence; these effects aggravated by H_2_O_2_-induced oxidative damage. Nevertheless, knockdown of NOX2 or overexpression of SIRT1 with adenovirus vector reduced progerin-associated premature senescence to maintain fenestrae through deacetylating p53. Furthermore, more Ac p53 K381 and progerin co-localized with accumulation of actin filament (F-actin) in the nuclear envelope of H_2_O_2_-treated HSECs; in contrast, these effects were rescued by overexpressing SIRT1. In conclusion, NOX2-dependent oxidative damage aggravates defenestration in HSECs via progerin-associated premature senescence; SIRT1-mediated deacetylation of p53 maintains fenestrae and attenuates liver fibrogenesis through inhibiting premature senescence.

## Introduction

Premature senescence, occurring when suffering from noxious stimuli, is characterized by inhibition of cell proliferation in advance and accumulation of damage, as well as involves in cellular dysfunction and various chronic diseases [1]. Emerging evidence confirms that the phenotypes and function of all intrahepatic cells have changed, particularly the loss of fenestration in hepatic sinusoidal endothelial cells (HSECs), due to older age; furthermore, the defenestration and capillarization in HSECs, as well as cirrhosis are observed in some premature senescence-related disease paradigms [2, 3]. These researches indicate that premature senescence in HSECs may be closely related to its defenestration and liver fibrogenesis. Hence, elucidation of the underlying mechanisms for premature senescence in HSECs may be a key to our understanding of defenestration and liver fibrosis pathogenesis.

The contraction and dilatation of fenestrae in HSECs and its differentiation are regulated by actin cytoskeleton (including F-actin) [4]. Our previous studies reveal that oxidative stress facilitates defenestration in HSECs via F-actin remodeling in liver fibrogenesis [5, 6]. Novel findings show that lamins and their associated proteins, which regulate nucleoskeleton and cytoskeleton, affect cellular differentiation and senescence [7, 8]. Especially, progerin is a mutant Lamin A protein, and its accumulation brings about abnormal nucleoskeleton and cell premature senescence to promote the occurrence and development of chronic liver diseases [7-9]. Thus, we speculate that progerin may contribute to premature senescence-associated defenestration in HSECs, via abnormal cytoskeleton remodeling.

Sirtuin 1 (SIRT1) is an important protector against oxidative stress and senescence to reverse the progression of chronic liver diseases [10]. Recent researches emphasize that overexpressing or activating SIRT1 inhibits hepatic senescence and activation of HSCs to ameliorate liver fibrosis [11, 12]. Besides, a major finding demonstrates that the activation of SIRT1 prevents endothelial cell from oxidative stress-induced senescence and dysfunction [13, 14]. Nevertheless, the effects of SIRT1 on premature senescence and defenestration in HSECs in liver fibrosis remain elusive.

Herein, our present study investigates the underlying mechanisms and the intervening target linking premature senescence and defenestration in HSECs, as well as the role of SIRT1 in defenestration *in vitro* and *in vivo*. We specifically focus on the SIRT1-mediated deacetylation that attenuates premature senescence-associated defenestration in HSECs in liver fibrosis.

## Results

### Premature senescence is induced by oxidative stress, with decrease of SIRT1 during defenestration in HSECs of CCl_4_-induced liver fibrogenesis

In our present study, fenestrae in hepatic sinusoidal endothelium disappeared completely on the 6^th^ day (Fig 1A), along with the high expression of α-SMA and vWF in CCl_4_-induced rat models on the 28^th^ day (Appendix Fig S1A-C, E); meanwhile, the serum ALT and AST levels increased, along with the augment of NOX2 protein level in primary rat HSECs which were isolated from CCl_4_-induced rat models (Appendix Fig S1D and E). These data indicated that CCl_4_ induced defenestration and capillarization in HSECs via oxidative stress. Interestingly, the senescence-associated β-galactosidase (SA-β-Gal)-positive cells increased in primary rat HSECs; moreover, the western blotting and immunofluorescence showed that a time-dependent elevation of the progerin protein expression in HSECs of CCl_4_-induced rat models implied that progerin might closely associated to premature senescence in CCl_4_-induced defenestrated HSECs (Fig 1B-D). However, the SIRT1 expression was down-regulated, with the enhance of the protein levels of Ac p53 K381 and total p53 in primary HSECs of CCl_4_-induced rat models (Fig 1C); the immunofluorescence and the immunohistochemical staining showed few expression of SIRT1 but much expression of Ac p53 K381 in vWF-positive hepatic sinusoidal endothelium (Fig 1E and F). Hence, these results confirmed that in the process of CCl_4_-induced defenestration in HSECs and liver fibrosis, oxidative stress triggered progerin-associated premature senescence, with decrease of SIRT1-mediated deacetylation.

**Figure 1.**
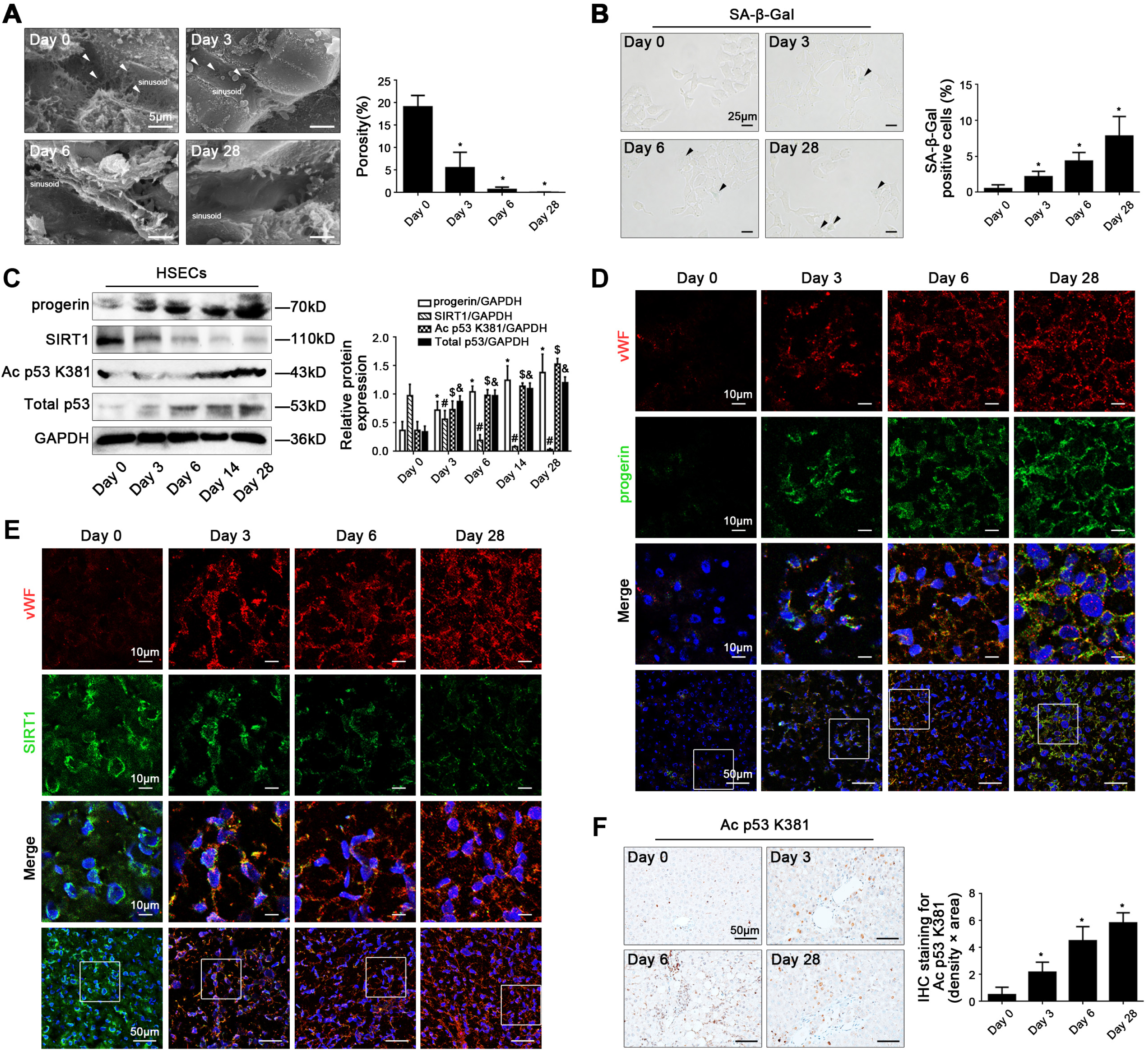
CCl_4_ induces progerin-associated premature senescence in defenestrated HSECs during liver fibrogenesis. (**A**) Magnification of scanning electron micrograph (SEM) of hepatic sinusoidal endothelium in CCl_4_-induced rat models (Day 0, Day 3, Day 6, and Day 28), revealing fenestrae structures in hepatic sinusoidal endothelium (Scale bar: 5 μm). The white triangles indicated fenestrae in hepatic sinusoidal endothelium. The porosity of hepatic sinusoidal endothelium was quantified in the graph, right. *P<0.05 versus Day 0. (**B**) The SA-β-Gal activity on primary HSECs, isolated from CCl_4_-induced rat models (Day 0, Day 3, Day 6, and Day 28), was observed by SA-β-Gal staining (Scale bar: 25 μm). The black triangles indicated the SA-β-Gal-positive cells. The SA-β-Gal-positive cells were quantified in the graph, right. *P<0.05 versus Day 0. (**C**) Representative immunoblots of progerin, SIRT1, Ac p53 K381, and total p53 of primary HSECs, isolated from CCl_4_-induced rat models (Day 0, Day 3, Day 6, Day 14, and Day 28). The relative protein expression was quantified in the graph, right. *P<0.05 versus progerin protein level on Day 0; ^#^P<0.05 versus SIRT1 protein level on Day 0; ^$^P<0.05 versus Ac p53 K381 protein level on Day 0; ^&^P<0.05 versus total p53 protein level on Day 0. (**D**) The immunofluorescent co-localization of vWF (red) with progerin (green) of liver biopsy specimens in CCl_4_-induced rat models (Day 0, Day 3, Day 6, and Day 28), visualized by confocal microscopy (Scale bar: 10 μm, 50 μm). Nuclear were showed by DAPI (blue). (**E**) The immunofluorescent co-localization of vWF (red) with SIRT1 (green) of liver biopsy specimens in CCl_4_-induced rat models (Day 0, Day 3, Day 6, and Day 28), visualized by confocal microscopy (Scale bar: 10 μm, 50 μm). Nuclear were showed by DAPI (blue). (**F**) The immumohistochemical (IHC) staining for Ac p53 K381 of liver biopsy specimens in CCl_4_-induced rat models (Day 0, Day 3, Day 6, and Day 28) (Scale bar: 50 μm). The semi-quantitative score of IHC staining for Ac p53 K381 was in the graph, right. *P<0.05 versus Day 0. n=6 per group.

### SIRT1 overexpression inhibits progerin-associated premature senescence to alleviate CCl_4_-induced defenestration in HSECs and liver fibrogenesis

To evaluate the role of SIRT1-mediated deacetylation in premature senescence and defenestration in HSECs *in vivo*, SIRT1 adenoviral vector was in advance transferred to CCl_4_-induced liver fibrogenesis rat models to ubiquitously overexpress SIRT1. There were about 70% of HSECs infected after injection. The H&E staining showed that SIRT1 adenoviral vector attenuated CCl_4_-induced liver acute injury and fibrogenesis on the 6^th^ and the 28^th^ day (Appendix Fig S2A). The vWF expression and the data of SEM demonstrated that overexpressing SIRT1 with adenoviral vector could maintain fenestrae and reduce capillarization in hepatic sinusoidal endothelium (Fig 2A and B; Appendix Fig S2B). Meanwhile, compared to the vehicle group, the NOX2 protein level, the H_2_O_2_ content, and mito-ROS increased in primary HSECs of the CCl_4_ group and the CCl_4_+AV-CTR group; these effects were reduced by the SIRT1 adenoviral vector (Fig 2A, C and D), indicated inhibition of oxidative stress via overexpressing SIRT1. Furthermore, compared to the vehicle group, the protein levels of Ac p53 K381, total p53, and progerin, as well as the SA-β-Gal-positive cells were elevated in primary HSECs of the CCl_4_ group and the CCl_4_+AV-CTR group on the 6^th^ and the 28^th^ day, which were inhibited by overexpressing SIRT1 (Fig 2A and E; Appendix Fig S2C). Besides, the immunofluorescence showed that compared with the vehicle group, the co-localization of Ac p53 K381 with F-actin highly expressed in the nuclear envelope of HSECs in the CCl_4_ group and the CCl_4_+AV-CTR group on the 6^th^ and 28^th^ day; in contrast, in the CCl_4_+AV-SIRT1 group on the 6^th^ and 28^th^ day, less Ac p53 K381 co-localized with F-actin; meanwhile F-actin distributed uniformly around the cell membrane of HSECs (Fig 2F), implied that overexpressing SIRT1 with adenoviral vector attenuated progerin-associated premature senescence and F-actin remodeling to maintain fenestrae in HSECs.

**Figure 2.**
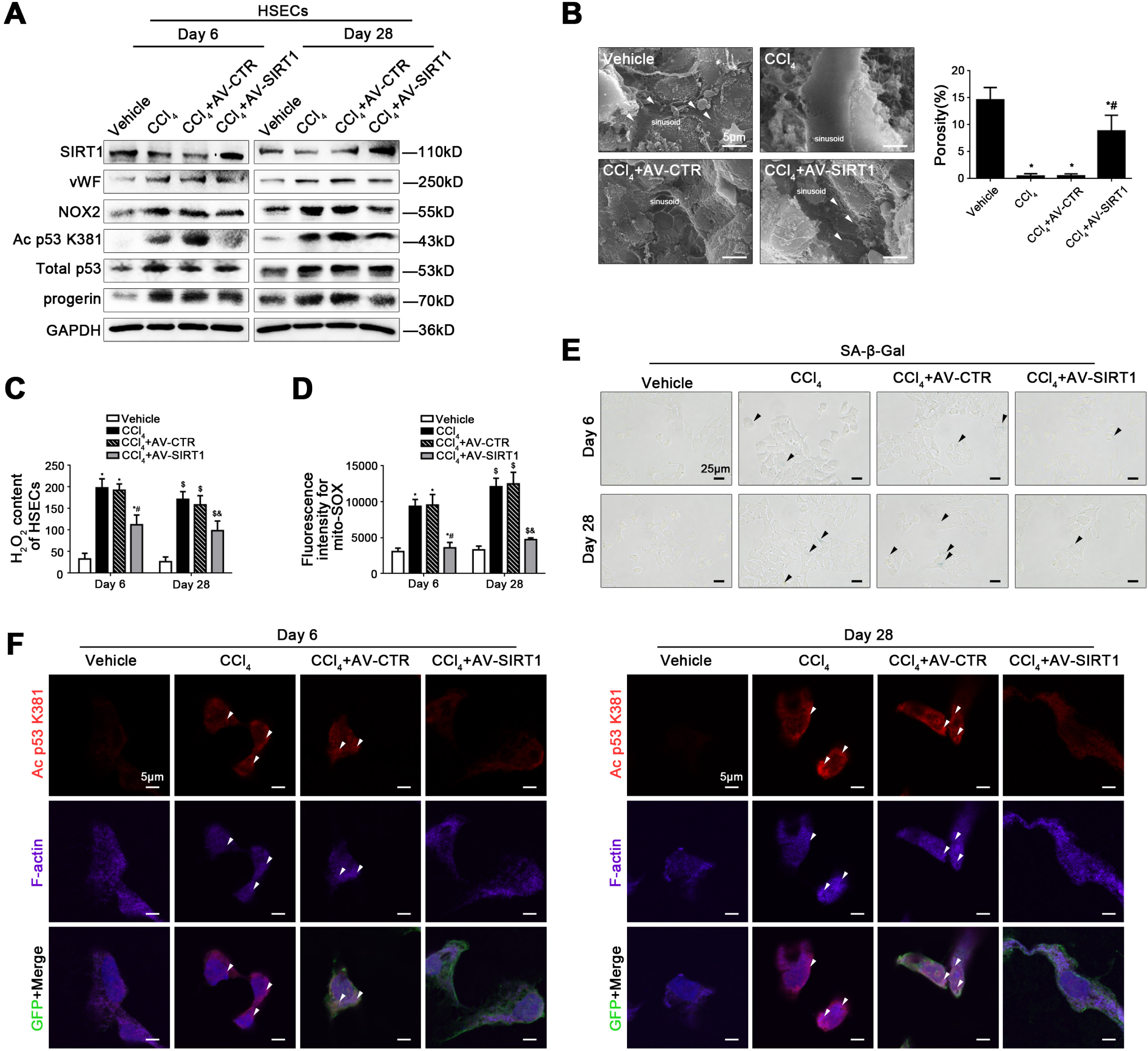
SIRT1 overexpression relieved progerin-associated premature senescence to attenuate CCl_4_-induced defenestration in HSECs. (**A**) Representative immunoblots of SIRT1, vWF, NOX2, Ac p53 K381, total p53, and progerin of primary HSECs, isolated from CCl_4_-induced rat models on Day 6 and Day 28. (**B**) Magnification of SEM of hepatic sinusoidal endothelium in CCl_4_-induced rat models on Day 6, revealing fenestrae structures in hepatic sinusoidal endothelium (Scale bar: 5 μm). The white triangles indicated fenestrae in hepatic sinusoidal endothelium. The porosity of hepatic sinusoidal endothelium was quantified in the graph, right. *P<0.05 versus the vehicle group; ^#^P<0.05 versus the CCl_4_ group and the CCl_4_+AV-CTR group. (**C**) The H_2_O_2_ content of primary HSECs, isolated from CCl_4_-induced rat models on Day 6 and Day 28. *P<0.05 versus the vehicle group on Day 6; ^#^P<0.05 versus the CCl_4_ group and the CCl_4_+AV-CTR group on Day 6; $P<0.05 versus the vehicle group on Day 28; ^&^P<0.05 versus the CCl_4_ group and the CCl_4_+AV-CTR group on Day 28. (**D**) Fluorescence intensity for mito-SOX of primary HSECs, isolated from CCl_4_-induced rat models on Day 6 and Day 28, measuring with flow cytometry. *P<0.05 versus the vehicle group on Day 6; ^#^P<0.05 versus the CCl_4_ group and the CCl_4_+AV-CTR group on Day 6; ^$^P<0.05 versus the vehicle group on Day 28; ^&^P<0.05 versus the CCl_4_ group and the CCl_4_+AV-CTR group on Day 28. (**E**) The SA-β-Gal activity on primary HSECs, isolated from CCl_4_-induced rat models on Day 6 and Day 28, was observed by SA-β-Gal staining (Scale bar: 25 μm). The black triangles indicated the SA-β-Gal-positive cells. (**F**) The immunocytochemical co-localization of Ac p53 K381 (red) with F-actin (purple) of primary HSECs, isolated from CCl_4_-induced rat models on Day 6 and Day 28, visualized by confocal microscopy (Scale bar: 5 μm). Adenovirus vectors were showed by GFP (green). Nuclear were showed by DAPI (blue). n=6 per group.

Taken together, these results demonstrated that activating SIRT1-mediated deacetylation relieved progerin-associated premature senescence and maintain cytoskeleton to attenuate CCl_4_-induced defenestration in hepatic sinusoidal endothelium and liver fibrogenesis.

### Progerin-associated premature senescence emerges in the process of defenestration in HSECs *in vitro*

*In vitro*, the fenestrae in freshly primary HSECs, which were isolated from normal rats and were cultured without growth factors for 5 days (Fig 3A), shrank rapidly from the 1^st^ day till the 3^rd^ day and disappeared completely on the 5^th^ day (Fig 3B). Interestingly, the SA-β-Gal-positive cells increased gradually with time (Fig 3C), along with the elevated protein levels of vWF, progerin, and Lamin A/C; whereas Lamin B1 protein expression was down-regulated (Fig 3D). These results indicated that the defenestration in HSECs was probably related to progerin-associated premature senescence.

**Figure 3.**
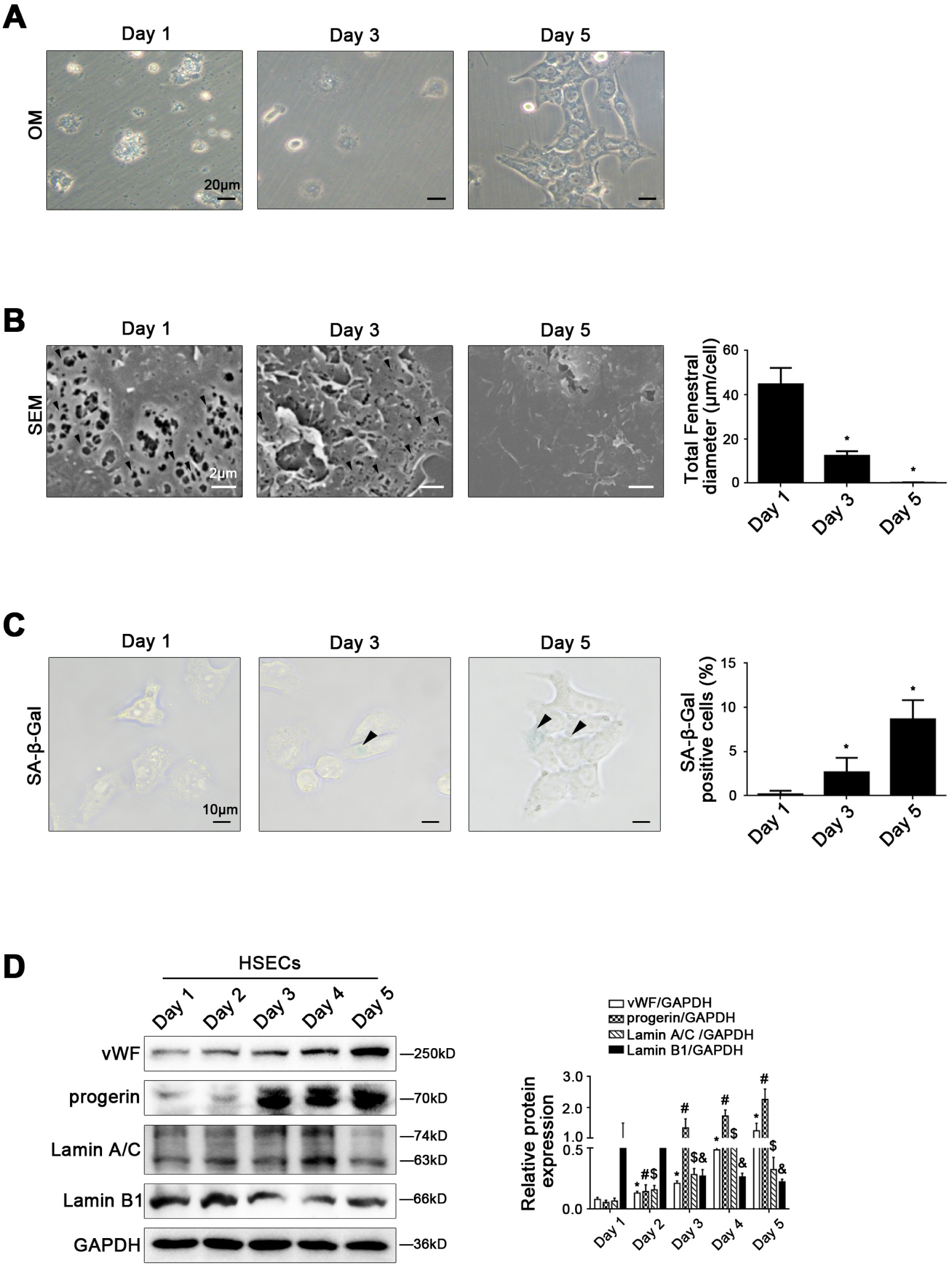
Progerin-associated premature senescence emerges in the process of defenestration in HSECs *in vitro*. Freshly primary HSECs, isolated from normal rats, were cultured without growth factors *in vitro* for 5 days. (**A**) The morphology of HSECs on Day 1, Day 3, and Day 5, observed by the microscope (Scale bar: 20 μm). (**B**) Magnification of SEM of HSECs on Day 1, Day 3, and Day5, revealing fenestrae structures in HSECs (Scale bar: 2 μm). The black triangles indicated fenestrae in HSECs. The total fenestral diameter was quantified in the graph, right. *P<0.05 versus Day 1. (**C**) The SA-β-Gal activity of HSECs on Day 1, Day 3, and Day 5, observed by SA-β-Gal staining (Scale bar: 10 μm). The black triangles indicated the SA-β-Gal-positive cells. The SA-β-Gal-positive cells were quantified in the graph, right. *P<0.05 versus Day 1. (**D**) Representative immunoblots of vWF, progerin, lamin A/C, and lamin B1 of primary HSECs. The relative protein expression was quantified in the graph, right. *P<0.05 versus vWF protein level on Day 1; ^#^P<0.05 versus progerin protein level on Day 1; $P<0.05 versus lamin A/C protein level on Day 1; ^&^P<0.05 versus lamin B1 protein level on Day 1.

### Oxidative stress inhibits SIRT1-mediated deacetylation and aggravates progerin-associated premature senescence to facilitate defenestration in HSECs

Freshly primary HSECs, isolated from normal rats, were stimulated with H_2_O_2_ at the different doses (0, 1.25, 2.5, 5, 10 μM) for 24 hours or at the dose (10 μM) from 12 hours to 48 hours. There was a time-dependent and a concentration-dependent up-regulation of NOX2, Ac p53 K381, and total p53 expression, with decrease of SIRT1 expression, in H_2_O_2_-treated primary rat HSECs (Fig 4A; Appendix Fig S3A). As expected, the data of SEM showed that the fenestrae in H_2_O_2_-treated HSECs disappeared on the 2^nd^ day, in advance of the control group (Fig 4B); meanwhile, the flow cytometry and immunocytochemistry showed that CD31, labeled continuous HSECs, highly express in H_2_O_2_-treated primary rat HSECs (Appendix Fig S3B and C), with the augment of vWF protein level (Fig 4A). Furthermore, the immunofluorescence showed that compared with the control group, the co-localization of NOX2 with F-actin highly expressed, accompany with accumulation of F-actin in the nuclear envelope of H_2_O_2_-treated HSECs on the 2^nd^ day (Fig 4C). These data suggested that H_2_O_2_-induced oxidative stress might inhibit SIRT1-mediated deacetylation of p53 and trigger F-actin remodeling to accelerate defenestration and capillarization in HSECs via NOX2.

**Figure 4.**
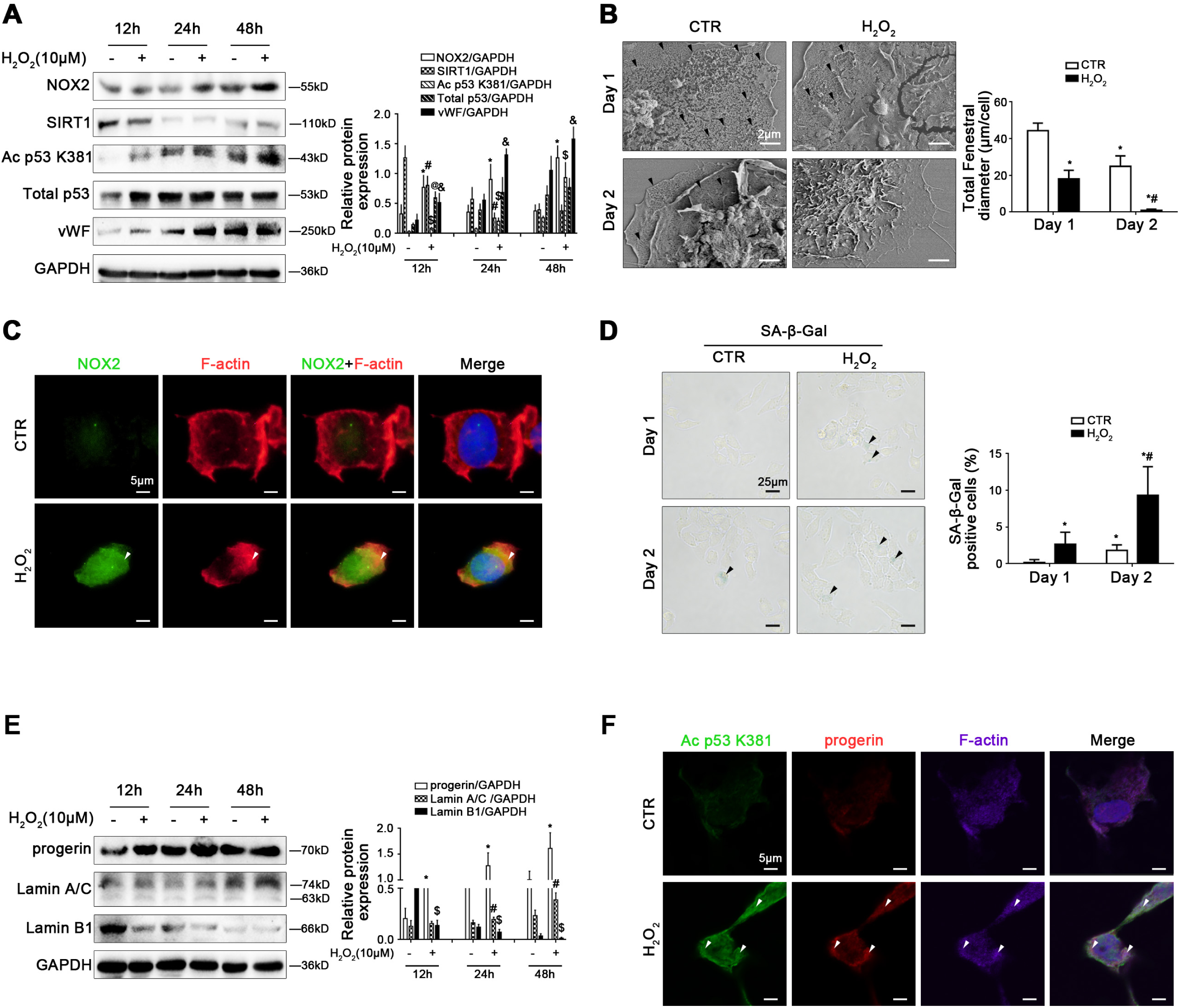
Oxidative stress inhibits SIRT1-mediated deacetylation and aggravates progerin-associated premature senescence to facilitate defenestration in HSECs. Freshly primary HSECs, isolated from normal rats and cultured *in vitro*, were treated with H_2_O_2_ (10 μM) from 12 hours to 48 hours. (**A**) Representative immunoblots of NOX2, SIRT1, Ac p53 K381, total p53, and vWF of HSECs in 12 hours, 24 hours, and 48 hours. The relative protein expression was quantified in the graph, right. *P<0.05 versus NOX2 protein level in the concurrent control group; ^#^P<0.05 versus SIRT1 protein level in the concurrent control group; ^$^P<0.05 versus Ac p53 K381 protein level in the concurrent control group; ^@^P<0.05 versus total p53 protein level in the concurrent control group; ^&^P<0.05 versus vWF protein level in the concurrent control group. (**B**) Magnification of SEM of HSECs in the CTR and H_2_O_2_ (10 μM) groups on Day 1 and Day 2, revealing the fenestrae structures (Scale bar: 2 μm). The black triangles indicated fenestrae in HSECs. The total fenestral diameter was quantified in the graph, right. *P<0.05 versus the CTR group on Day 1; ^#^P<0.05 versus the CTR group on Day 2. (**C**) The immunocytochemical co-localization of NOX2 (green) with F-actin (red) in primary rat HSECs on Day 2, visualized by confocal microscopy (Scale bar: 5 μm). Nuclear were showed by DAPI (blue). (**D**) The SA-β-Gal activity in primary rat HSECs on Day 1 and Day 2, was observed by SA-β-Gal staining (Scale bar: 25 μm). The black triangles indicated the SA-β-Gal-positive cells. The SA-β-Gal-positive cells were quantified in the graph, right. *P<0.05 versus the CTR group on Day 1; ^#^P<0.05 versus the CTR group on Day 2. (**E**) Representative immunoblots of progerin, Lamin A/C, and Lamin B1 of HSECs in 12 hours, 24 hours, and 48 hours. The relative protein expression was quantified in the graph, right. *P<0.05 versus progerin protein level in the concurrent control group; #P<0.05 versus Lamin A/C protein level in the concurrent control group; ^$^P<0.05 versus Lamin B1 protein level in the concurrent control group. (**F**) The immunocytochemical co-localization of Ac p53 K381 (green) with progerin (red) and F-actin (purple) of primary rat HSECs on Day 2, visualized by confocal microscopy (Scale bar: 5 μm). Nuclear were showed by DAPI (blue).

Besides, the SA-β-Gal staining, as well as the protein levels of progerin, Lamin A/C and Lamin B1 showed that H_2_O_2_-induced oxidative stress promoted progerin-associated premature senescence in HSECs, with decrease of Lamin B1 expression (Fig 4D and E). Compared to the control group, more progerin and Ac p53 K381 also were co-localized with F-actin in the nuclear envelope of H_2_O_2_-treated HSECs on the 2^nd^ day (Fig 4F). Hence, these results indicated that H_2_O_2_-induced oxidative stress triggered progerin-associated premature senescence through acetylation of p53 at lysine 381, and subsequently contributed to F-actin remodeling to aggravate defenestration in HSECs.

In addition, primary rat HSECs were transfected with p53 siRNA, progerin siRNA, or nontarget siRNA (called NC), and then administered with H_2_O_2_ (10 μM) for two days. We found that H_2_O_2_-induced strengthened activity of SA-β-Gal, was significantly reduced by silencing p53 with p53 siRNA (Appendix Fig S4A and B), suggested p53-mediated premature senescence in H_2_O_2_-treated HSECs. As expected, the data of SEM showed that silencing progerin with progerin siRNA attenuated H_2_O_2_-induced HSECs defenestration on the 2^nd^ day (Appendix Fig S4C and D), implied that inhibiting progerin-associated premature senescence could maintain HSECs fenestrae.

In consequence, H_2_O_2_-induced oxidative stress inhibited SIRT1-mediated deacetylation of p53 and triggered progerin-associated premature, and then brought about abnormal cytoskeleton remodeling to aggravate defenestration in HSECs.

### Inhibiting NOX2-dependent oxidative stress reduces progerin-associated premature senescence to maintain fenestrae in HSECs

To further delineate the molecular mechanism of oxidative stress-induced premature senescence and defenestration in HSECs, primary rat HSECs, isolated from normal rats and cultured *in vitro*, were transfected with NOX2 siRNA or nontarget siRNA (called NC), and then administered with H_2_O_2_ (10 μM) for two days. The transfection efficiency achieved 75%. As we expected, compared to the control group and the NC group, H_2_O_2_ caused the increase of mito-ROS, as well as the NOX2 mRNA and its protein high expression, as a result of NOX2-dependent oxidative stress, along with decrease of SIRT1 expression but increase of Ac p53 K381 expression; in contrast, these effects remarkably were inhibited by knockdown of NOX2 with NOX2 siRNA (Fig 5A-C). Moreover, the protein levels of progerin and Lamin A/C, as well as the activity of SA-β-Gal were elevated in HSECs, along with decrease of Lamin B1 expression; on the contrary, knockdown of NOX2 down-regulated the protein levels of progerin and Lamin A/C, but up-regulated the Lamin B1 level (Fig 5D and E), implied that inhibiting NOX2-dependent oxidation attenuated progerin-associated premature senescence. Additionally, the immunofluorescence and the SEM showed that accumulation of F-actin in the nuclear envelope of H_2_O_2_-treated HSECs, as well as its defenestration were triggered by oxidative stress, which were rescued by knockdown of NOX2 (Fig 5F and G). In short, inhibition of NOX2-dependent oxidative stress alleviates progerin-associated premature senescence to maintain fenestrae in HSECs.

**Figure 5.**
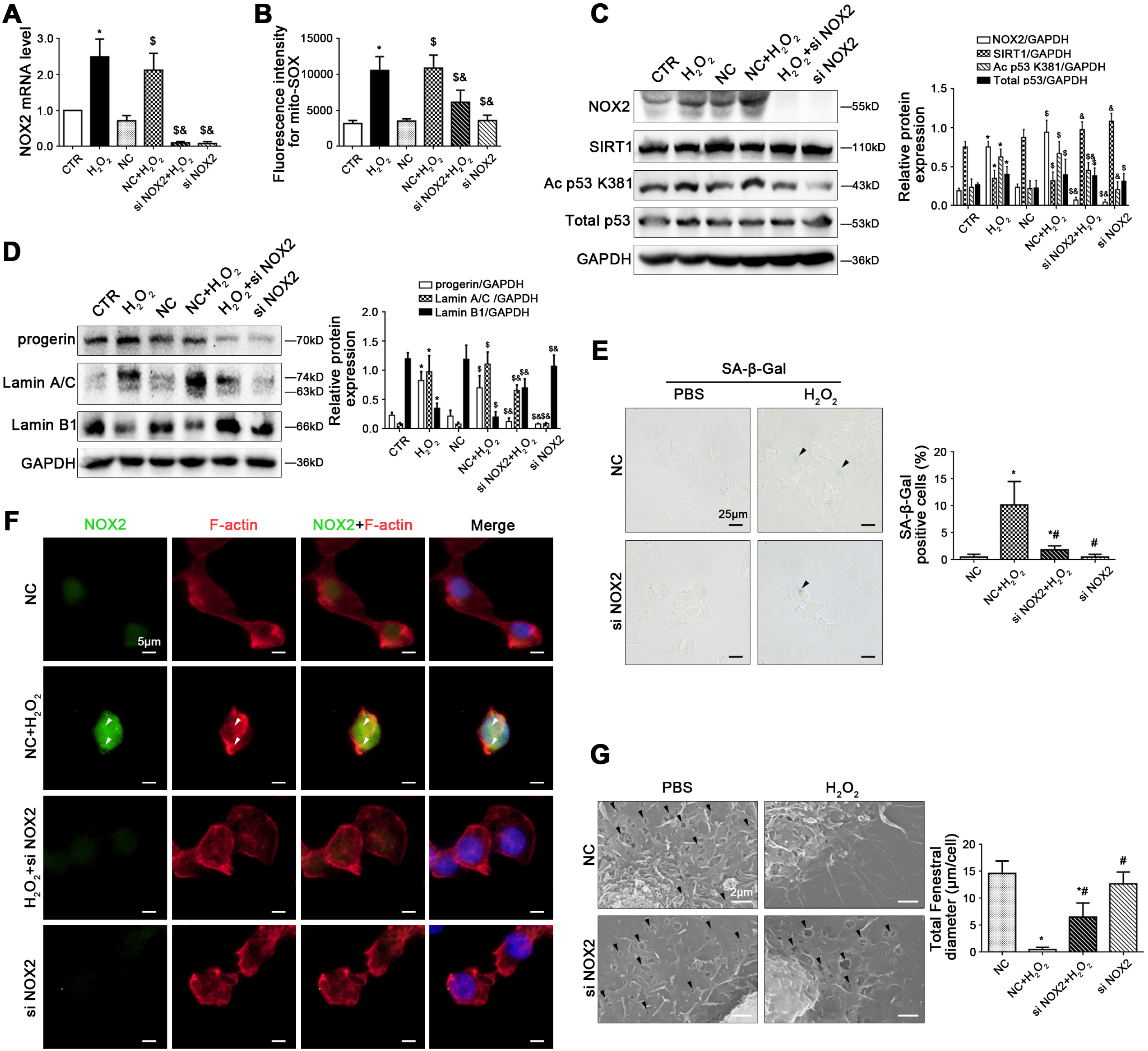
Inhibiting NOX2-dependent oxidative stress reduces progerin-associated premature senescence to maintain fenestrae in HSECs. Freshly primary rat HSECs, isolated from normal rats and cultured *in vitro*, were transfected with NOX2 siRNA or nontarget siRNA (called NC), and then administered with H_2_O_2_ (10 μM) for two days. (**A**) Real-time PCR analysis of NOX2 mRNA level in HSECs on Day 2. *P<0.05 versus the CTR group; ^$^P<0.05 versus the NC group; ^&^P<0.05 versus the NC+H_2_O_2_ group. (**B**) Fluorescence intensity for mito-SOX of HSECs on Day 2, measuring with flow cytometry. *P<0.05 versus the CTR group; ^$^P<0.05 versus the NC group; ^&^P<0.05 versus the NC+H_2_O_2_ group. (**C**) Representative immunoblots of NOX2, SIRT1, Ac p53 K381, and total p53 of HSECs on Day 2. The relative protein expression was quantified in the graph, right. *P<0.05 versus the CTR group; ^$^P<0.05 versus the NC group; ^&^P<0.05 versus the NC+H_2_O_2_ group. (**D**) Representative immunoblots of progerin, Lamin A/C, and Lamin B1 of HSECs on Day 2. The relative protein expression was quantified in the graph, right. *P<0.05 versus the CTR group; ^$^P<0.05 versus the NC group; ^&^P<0.05 versus the NC+H_2_O_2_ group. (**E**) The SA-β-Gal activity in HSECs on Day 2 in the four groups (NC, NC+H_2_O_2_, H_2_O_2_+si NOX2, si NOX2), was observed by the SA-β-Gal staining (Scale bar: 25 μm). The black triangles indicated the SA-β-Gal-positive cells. The SA-β-Gal-positive cells were quantified in the graph, right. *P<0.05 versus the NC group; ^#^P<0.05 versus the NC+H_2_O_2_ group. (**F**) The immunocytochemical co-localization of NOX2 (green) with F-actin (red) in primary HSECs on Day 2, visualized by confocal microscopy (Scale bar: 5 μm). Nuclear were showed by DAPI (blue). (**G**) Magnification of SEM of HSECs in the four groups (NC, NC+H_2_O_2_, H_2_O_2_+si NOX2, si NOX2), revealing fenestrae structures in HSECs (Scale bar: 2 μm). The black triangles indicated fenestrae in HSECs. The total fenestral diameter was quantified in the graph, right. *P<0.05 versus the NC group; ^#^P<0.05 versus the NC+H_2_O_2_ group.

### SIRT1-mediated deacetylation relieves NOX2-dependent oxidative stress to maintain fenestrae in HSECs via attenuating progerin-associated premature senescence

In order to characterize fenestrae in HSECs responding to SIRT1-mediated deacetylation in depth, primary rat HSECs, isolated from normal rats, were transfected with the SIRT1 adenovirus vector to overexpress SIRT1 (namely AV-SIRT1) or nontarget adenovirus vector (called AV-CTR), and then stimulated with H_2_O_2_ (10 μM) for two days. The protein levels of vWF, NOX2, and NOX4, as well as mito-ROS were enhanced by H_2_O_2_, which were decreased by overexpression of SIRT1 with adenovirus vector (Fig 6A and B), indicated that SIRT1 relieved H_2_O_2_-induced NOX2-dependent oxidative stress and mitochondrial dysfunction; meanwhile, compared with the control group, H_2_O_2_ caused more co-localization of NOX2 with F-actin in the nuclear envelope on the 2^nd^ day; whereas these effects were suppressed by SIRT1 adenovirus vector (Fig 6C), implied that SIRT1 reduced F-actin remodeling through inhibiting NOX2-dependent oxidation.

**Figure 6.**
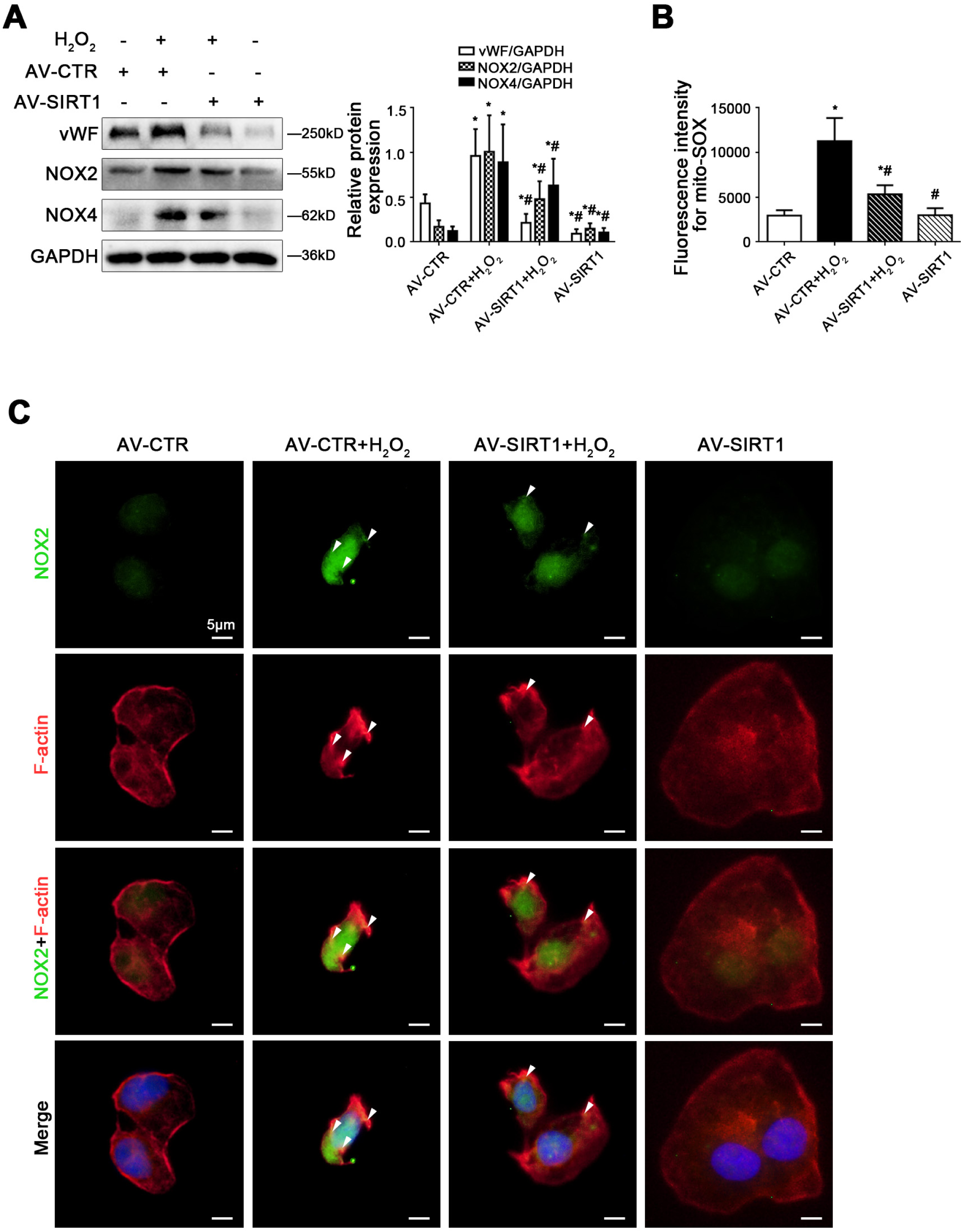
SIRT1 reduced F-actin remodeling through inhibiting NOX2-dependent oxidative stress. Freshly primary rat HSECs, isolated from normal rats and cultured *in vitro*, were transfected with the SIRT1 adenovirus vector to overexpress SIRT1 (called AV-SIRT1) or nontarget adenovirus vector (called AV-CTR), and then treated with H_2_O_2_ (10 μM) for two days. (**A**) Representative immunoblots of vWF, NOX2, and NOX4 of primary HSECs on Day 2 in four groups (AV-CTR, AV-CTR+H_2_O_2_, AV-SIRT1+H_2_O_2_, AV-SIRT1). The relative protein expression was quantified in the graph, right. *P<0.05 versus the AV-CTR group; ^#^P<0.05 versus the AV-CTR+H_2_O_2_ group. (**B**) Fluorescence intensity for mito-SOX of primary HSECs on Day 2, measuring with flow cytometry. *P<0.05 versus the AV-CTR group; ^#^P<0.05 versus the AV-CTR+H_2_O_2_ group. (**C**) The immunocytochemical co-localization of NOX2 (green) with F-actin (red) of primary HSECs on Day 2, visualized by confocal microscopy (Scale bar: 5 μm). Nuclear were showed by DAPI (blue).

Furthermore, the SA-β-Gal staining showed that overexpressing SIRT1 with the adenovirus vector reduced H_2_O_2_-induced senescence (Fig 7A). The protein levels of SIRT1, Ac p53 K381, total p53, progerin, Lamin A/C, and Lamin B1 in nuclei and cytoplasm showed that compared to the control group, the expression of Ac p53 K381, progerin, and Lamin A/C were strongly elevated, with the decrease of SIRT1 and Lamin B1 in nuclei of H_2_O_2_-treated HSECs; in the contrary, these effects were recused by overexpression of SIRT1 with adenovirus vector (Fig 7B), indicated that SIRT1 overexpression suppressed progerin-associated premature senescence. The co-immunoprecipitation (Co-IP) assay revealed that the co-precipitation of Ac p53 K381 with progerin was enhanced in nuclei of H_2_O_2_-treated HSECs, whose interaction was disrupted by SIRT1 adenovirus vector (Fig 7C); meanwhile, the immunofluorescence showed more Ac p53 K381 and progerin co-localized with F-actin in the nuclear envelope of H_2_O_2_-treated HSECs on the 2^nd^ day; whereas less co-localization of Ac p53 K381 and progerin with F-actin were displayed in the AV-SIRT1 group and the AV-SIRT1+H_2_O_2_ group (Fig 7D). These data demonstrated that overexpressing SIRT1 blocked the nuclei Ac p53 K381-progerin interaction. In addition, the data of SEM revealed that overexpression of SIRT1 with adenovirus vector rescued H_2_O_2_-induced defenestration in HSECs (Fig 7E).

**Figure 7.**
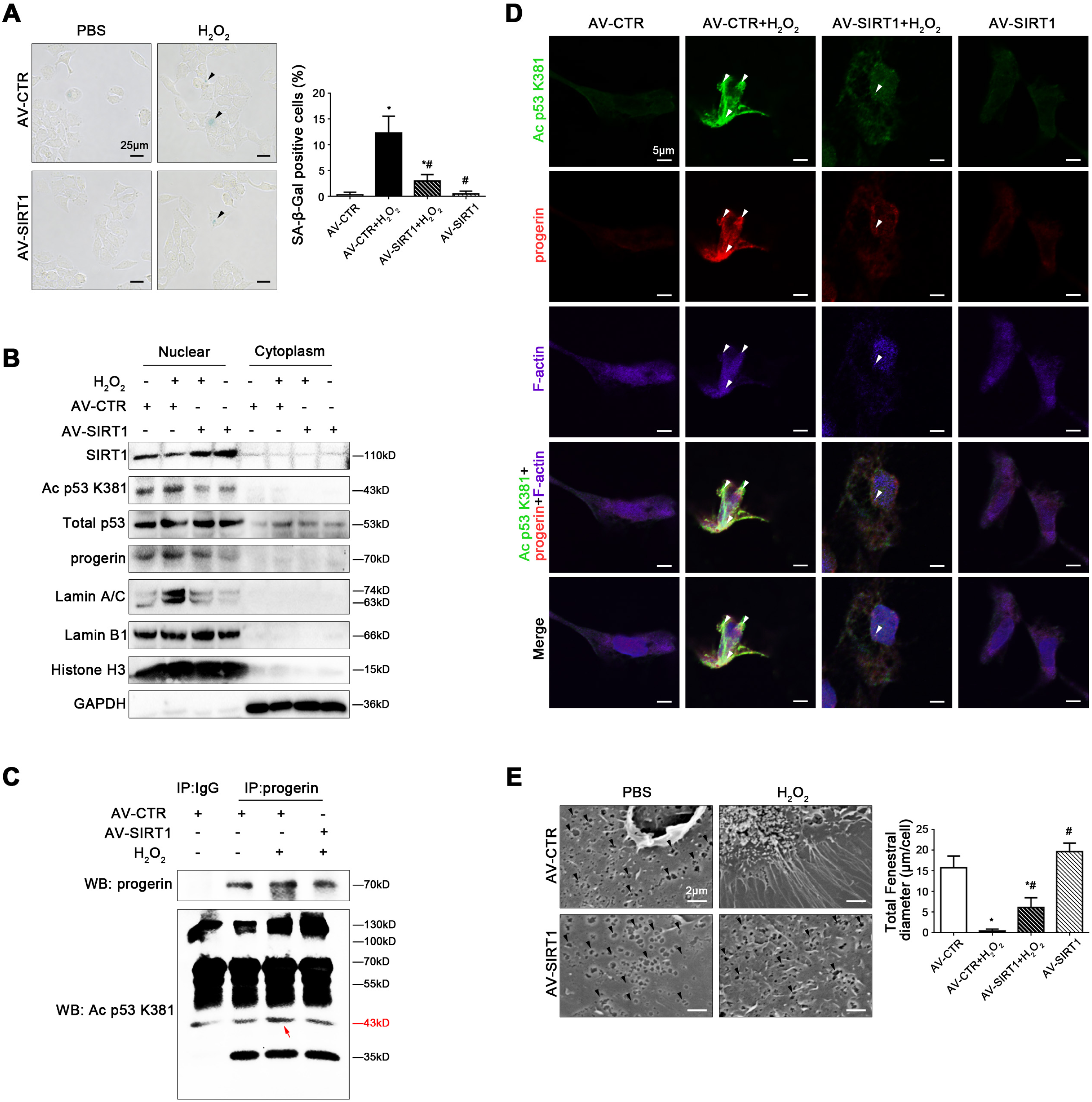
SIRT1-mediated deacetylation maintains fenestrae in HSECs via inhibiting progerin-associated premature senescence. Freshly primary rat HSECs, isolated from normal rats and cultured *in vitro*, were transfected with the SIRT1 adenovirus vector to overexpress SIRT1 (called AV-SIRT1) or nontarget adenovirus vector (called AV-CTR), and then treated with H_2_O_2_ (10 μM) for two days. (A) The SA-β-Gal activity in HSECs on Day 2 in the four groups (AV-CTR, AV-CTR+H_2_O_2_, H_2_O_2_+AV-SIRT1, AV-SIRT1), was observed by SA-β-Gal staining (Scale bar: 25 μm). The black triangles indicated the SA-β-Gal-positive cells. The SA-β-Gal-positive cells were quantified in the graph, right. *P<0.05 versus the AV-CTR group; ^#^P<0.05 versus the AV-CTR+H_2_O_2_ group. (B) Primary rat HSECs were extracted nuclear and cytoplasmic protein and were detected their protein levels. Representative immunoblots of SIRT1, Ac p53 K381, total p53, progerin, Lamin A/C, and Lamin B1 in nuclei and cytoplasm of HSECs in four groups (AV-CTR, AV-CTR+H_2_O_2_, AV-SIRT1+H_2_O_2_, AV-SIRT1). (**C**) Interaction of progerin with Ac p53 K381 was detected by the co-IP assay. Progerin of HSECs was individually immunoprecipitated, as well as Ac p53 K381 and progerin subjected to immunoblotting analysis as indicated. (**D**) The immunocytochemical co-localization of Ac p53 K381 (green) with progerin (red) and F-actin (purple) in primary HSECs on Day 2 in four groups (AV-CTR, AV-CTR+H_2_O_2_, AV-SIRT1+H_2_O_2_, AV-SIRT1), visualized by confocal microscopy (Scale bar: 5 μm). Nuclear were showed by DAPI (blue). (**E**) Magnification of SEM of HSECs in the four groups (AV-CTR, AV-CTR+H_2_O_2_, AV-SIRT1+H_2_O_2_, AV-SIRT1), revealing fenestrae structures in HSECs (Scale bar: 2 μm). The black triangles indicated fenestrae in HSECs. The total fenestral diameter was quantified in the graph, right. *P<0.05 versus the AV-CTR group; ^#^P<0.05 versus the AV-CTR+H_2_O_2_ group.

Therefore, these results confirmed that SIRT1-mediated deacetylation of p53 relieved NOX2-dependent oxidative stress and inhibited progerin-associated premature senescence to maintain fenestrae in HSECs.

## Discussion

Our present study discovers that NOX2-dependent oxidative stress triggers progerin-associated premature senescence through inhibiting SIRT1-mediated deacetylation of p53, and subsequently aggravates cytoskeleton remodeling to promote defenestration in HSECs; overexpressing SIRT1 with adenovirus vector strongly activates deacetylation of p53, and then relieves progerin-related premature senescence to maintain fenestrae in HSECs and reverse liver fibrogenesis (Fig 8).

**Figure 8.**
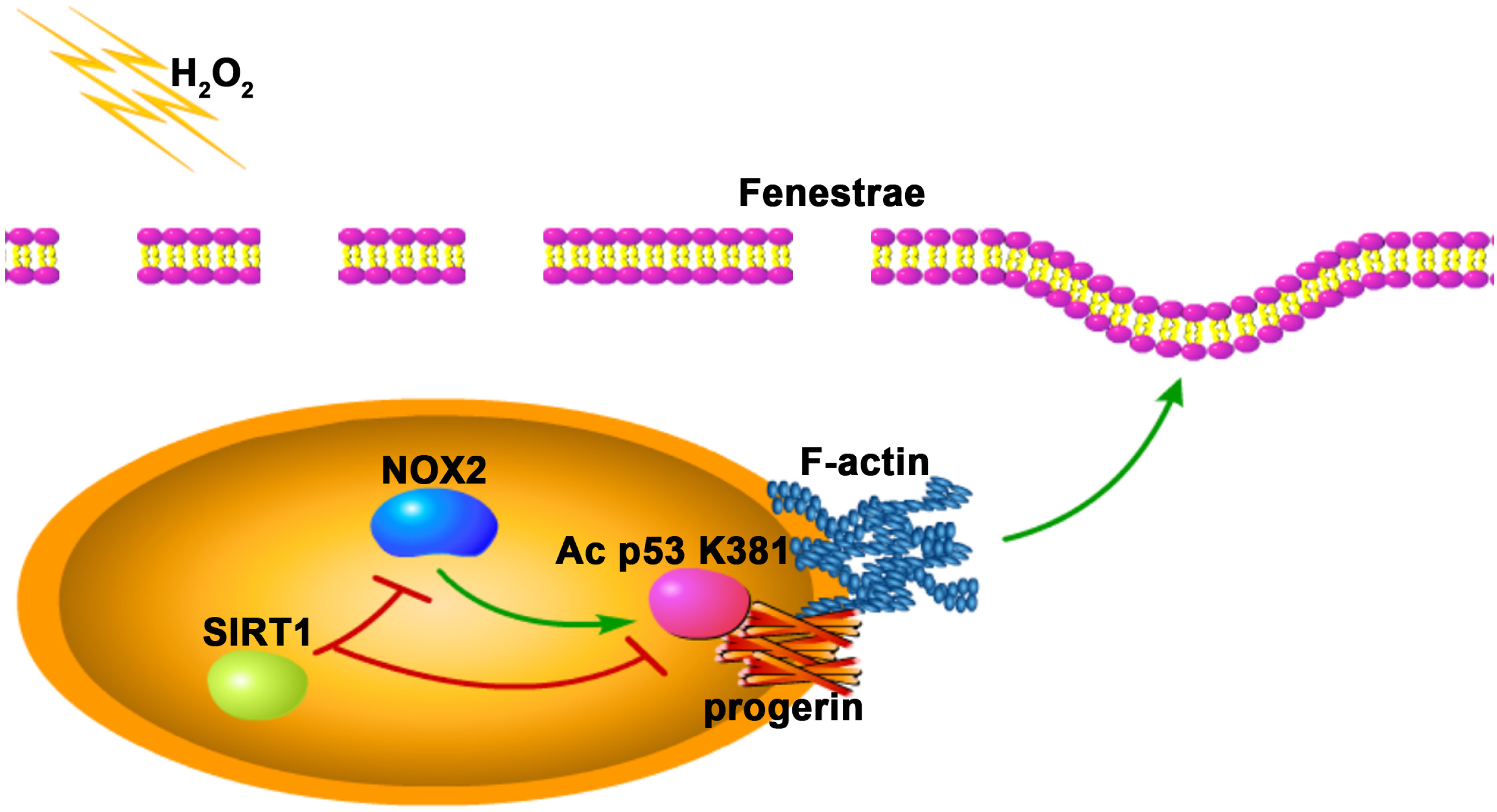
A schematic view of major signal transduction pathways, involves in the conclusion that SIRT1-mediated deacetylation of p53 relieved NOX2-dependent oxidative stress and inhibited progerin-associated premature senescence to maintain fenestrae in HSECs.

There is mounting evidence that premature senescence, induced by harmful stimuli, is a permanently deteriorated process about cell cycle arrest in advance, which involves in endothelial cells dysfunction and its related diseases as diverse as diabetes, lipodystrophy, and atherosclerosis, to name a few [15, 16]. In our previous findings, premature senescence emerges in liver tissue due to acute liver injury and liver fibrogensis [17]. In addition, in older animal models and premature aging-related disease cases, abnormal differentiation and dysfunction in HSECs is attributed to premature senescence [2, 3]. In this study, we similarly found that premature senescence in HSECs occurred in the process of fenestrae disappearance *in vitro*; oxidative stress enhanced premature senescence to aggravate defenestration in H_2_O_2_-treated HSECs and in HSECs of CCl_4_-induced fibrogenesis. Nevertheless, the underlying mechanism about how oxidative stress-induced premature senescence triggers defenestration in HSECs, is still elusive.

Lamins and its associated protein, especially Lamin A and Lamin B1, forms an interface with nuclear membrane and nuclear pore complexes [18], while alteration of lamins involves in cellular premature senescence and age-associated diseases. Interestingly, some literatures regarding the influence of lamins on premature senescence and liver diseases are controversial. For instance, the disruption or mutantion of lamins led to hepatocytes abnormal differentiation and premature aging to induce steatohepatitis [7]. Progerin, a mutant of Lamin A, is deemed to a moderator for premature aging and dysfunction in endothelial cells [19]. The depletion of Lamin B contributes to chronic inflammation [20]. Our intriguing discoveries revealed that premature senescence in HSECs, along with abnormal accumulation of progerin and the decrease of Lamin B1, during H_2_O_2_-treated or CCl_4_-induced HSECs defenestration; whereas knockdown of progerin could reduce premature senescence to recover fenestrae in H_2_O_2_-treated HSECs. These findings indicate that A-type and B-type lamins, are required for HSECs fenestrae; meanwhile, oxidative stress induced-premature senescence leads to defenestration in HSECs via targeting progerin.

However, how progerin-associated premature senescence, induced by oxidative stress, aggravates defenestration in HSECs? NOX2 activation and its derived ROS are implicated in oxidative damage of vascular function in aging-related diseases [21]; moreover, activating NOX2 accelerates liver fibrosis during aging [22]. Our findings showed that H_2_O_2_ elevated the NOX2 protein level and mitochondrial dysfunction to trigger progerin-associated premature senescence, leading to defenestration in HSECs; silencing NOX2 could rescue mitochondrial function and premature senescence, and subsequently maintain HSECs fenestrae. These data demonstrated that premature senescence and defenestration in HSECs, which induced by oxidative stress, attributed to NOX2 activation. Besides, emerging reports show that lamins play a critical role in nucleoskeleton and cytoskeleton; whereas, accumulation of progerin contributes to perturbations in actin organization [19], implying that progerin-associated premature senescence modulates cytoskeleton to affect cell phenotype. Some novel studies report that lamins-related aging is regulated by transcription factors (such as p53, NF-κB, etc.) [23, 24]. In our present study, more co-localization of NOX2 and F-actin displayed in the nuclear envelope of H_2_O_2_-treated HSECs, whose fenestrae disappeared; on the contrary, knockdown of NOX2, which reduced oxidative stress-induced premature senescence, inhibited F-actin remodeling to rescue defenestration in HSECs. In brief, NOX2-dependent oxidative stress triggered progerin-associated premature senescence to result in defenestration in HSECs via F-actin remodeling.

SIRT1, a class III histone deacetylase, serves as an important modulator of metabolism, cellular survival, and lifespan [10]. Some recent evidence reports that the activation of SIRT1 confers protective effects on hepatic senescence and HSCs activation to ameliorate liver fibrosis [11, 12]. The previous investigations reveal that SIRT1 protects vascular endothelial cells against age-related endothelial dysfunction; whereas few researches explore the role of SIRT1 in morphology and function of HSECs. In our present study, oxidative damage suppressed SIRT1-medatied deacetylation of p53 in H_2_O_2_-treated HSECs; in paralleling, the Ac p53 K381 protein expression highly expressed in nuclei and co-localized with more progerin and F-actin in the nuclear envelope of H_2_O_2_-treated HSECs. In contrast, overexpression of SIRT1 with adenovirus vector inhibited NOX2-dependent oxidative stress, relieved premature senescence, and blocked abnormal accumulation of progerin and F-actin remodeling, to attenuate H_2_O_2_-treated and CCl_4_-induced defenestration in HSECs and liver fibrogenesis. It follows that activation of SIRT1 effectively protects against defenestration in HSECs through inhibiting progerin-associated premature senescence.

Although this study provides new insights into the mechanisms of premature senescence in HSECs defenestration, there are still some limitations to our present study. The mechanism about reduction of Lamin B1 in HSECs defenestration needs to lucubrate. Besides, how progerin-associated premature senescence affects liver sinusoid capillarization remains elusive.

In conclusion, NOX2-dependent oxidative damage aggravates defenestration in HSECs, as a result of progerin-associated premature senescence; SIRT1-mediated deacetylation of p53 maintains fenestrae in HSECs and attenuates liver fibrogenesis via inhibition of progerin-related premature senescence.

## Materials and Methods

### Reagents and antibodies

The reagents used included carbon tetrachloride (Sigma-Aldrich, 56-23-5), 30%H_2_O_2_ (Hydrogen peroxide 30%, Sigma-Aldrich, 1.07298), DAPI (Sigma-Aldrich, D9542), Alexa Fluor(tm) 647 Phalloidin (Thermo, A22287), protease cocktails inhibitor (Beyotime, P1005), and PMSF (Phenylmethanesulfonyl fluoride, Beyotime, ST506).

The primary antibodies included anti-α-SMA (Boster, BM0002), anti-vWF (Santa Cruz, SC-365712), anti-CD32b (ZEN-bioscience, 382560), anti-NOX2 (Proteintech, 19013-1-AP), anti-NOX4 (Proteintech, 14347-1-AP), anti-Lamin A/C (Cell Signaling Technology, 4777S), anti-Lamin B1 (Proteintech, 66095-1-Ig), anti-progerin (Santa Cruz, sc-81611), anti-p53 (Abcam, ab131442), anti-p53 (acetyl K381) (Abcam, ab61241), anti-SIRT1 (Abcam, ab110304), anti-Histone H3 (Proteintech, 17168-1-AP), and anti-GAPDH (Proteintech, 60004-1). HRP-conjugated Affinipure Goat Anti-Mouse IgG(H+L) (Proteintech, SA00001-1), HRP-conjugated Affinipure Goat Anti-Rabbit IgG(H+L) (Proteintech, SA00001-2), FITC-labeled goat anti-rabbit IgG(H+L) (Beyotime, a0562), and Cy3-labeled goat anti-mouse IgG (H+L) (Beyotime, a0521) were used for secondary antibodies.

### Animal experimental design

Sprague-Dawley (SD) rats were provided by the Laboratory Animal Center (Henan University of Chinese Medicine, China) and were approved by the Committee on the Ethics of Animal Experiments of Southern Medical University. Animals were housed under a 12:12 h light/dark cycle at 22–24 °C.

#### Establishment of CCl_4_-induced liver fibrogenesis rat models

Normal male SD rats (180-220g) were subjected to intraperitoneal injection of 40% carbon tetrachloride (CCl_4_)-olive oil solution at 2ml/kg body weight, twice a week for 28 days. On Day 0, 3, 6, and 28, CCl_4_-induced rat models were randomly sacrificed (n = 6 per group).

#### The treatment of SIRT1 adenovirus vector

To investigate the role of SIRT1 in fenestrae in primary HSECs and liver fibrogenesis, the GFP-SIRT1-adenovirus vector and the GFP-blank vector were produced by Hanbio AdenoVector Institute (Shanghai, China) and the dose of 10^11^ viral particles was injected through caudal vein to rats one week before the intraperitoneal injection of CCl_4_-olive oil solution. We employed the CCl_4_-induced liver fibrosis rat models (n = 6 per group for 6 days and n = 6 per group for 28 days). The vehicle group (n = 6 per group for 6 days and n = 6 per group for 28 days) was subjected to intraperitoneal injection of the same volume of olive oil, twice a week for 28 days. The AV-CTR+CCl_4_ group and the AV-SIRT1+CCl_4_ group (n = 6 per group for 6 days and n = 6 per group for 28 days) was subjected to intraperitoneal injection of CCl_4_-olive oil solution twice a week after administering vectors. On Day 6 and 28, the rat models were randomly sacrificed. The SIRT1 sequences were used: sense (5-CGGGCCCTCTAGACTCGAGCGGCCGCATGATTGGCACCGATCCTC-3).

### Histological analysis and immunohistochemistry

Paraffin sections (4 μm) of liver tissue of the model rats were prepared with hematoxylin and eosin (H&E) staining. The rat liver histological inflammation and fibrosis stage are assessed with the Ishak inflammation and fibrosis score (from Supplementary method). Immunohistochemical detection of α-SMA, vWF, and Ac p53 K381, were performed on paraffin sections (3 μm) of liver tissue, and subsequent sections were exposed to HRP-antibody colored with DAB, and visualized by microscopy (BX51, Olympus, Japan). The degree of liver fibrosis and the number of α-SMA-, vWF-, or Ac p53 K381-positive cells were quantified with Image J software.

### Immunofluorescence staining

Paraffin sections (3 μm) of liver tissue of the model rats were prepared for immunofluorescence, incubated with primary antibody overnight, followed by the secondary antibody, and then mounted with DAPI. The primary antibodies included anti-progerin (1:50), anti-SIRT1 (1:200), and anti-vWF (1:200). The secondary antibodies included FITC-labeled goat anti-rabbit IgG (H+L) (1:200) and Cy3-labeled goat anti-mouse IgG (H+L) (1:200).

### Cell isolation, identification, culture and treatment

Primary rat HSECs were isolated from normal male SD rats and identified by SEM, based on the modified method [5, 6]. Primary rat HSECs were cultured in plates with medium comprising 80% MCDB131 (Gibco, 10372019) and 20% fetal bovine serum (FBS, Biological Industries, 04-007-1A). Primary rat HSECs HSECs were stimulated by H_2_O_2_ with a concentration gradient of 0, 1.25, 2.5, 5, 10 μM for 24 hours, or a time gradient of 0, 12, 24, 36, 48 hours.

### Measurement of the activity of senescence-associated β-galactosidase (SA-β-Gal)

The activity of SA-β-Gal in primary rat HSECs was determined using 5-bromo-4-chloro-3-indolyl P3-D-galactoside (X-gal), according to the manufacturer instruction (Senescence-associated β-galactosidase Staining Kit, Beyotime, C0602). SA-β-Gal-positive cells (blue color) were counted under microscope.

### Scanning electron microscopy (SEM)

The liver tissue of the model rats and primary rat HSECs were fixed with 2.5% glutaldehyde and subsequently dehydrated and then coated with gold using the coating apparatus, based on the modified method [5]. Eventually, fenestrae in primary rat HSECs of samples were observed with SEM at 15-kV acceleration voltage.

### SIRT1 adenovirus transfection

The recombinant adenovirus was produced by Hanbio AdenoVector Institute (Shanghai, China). To construct Flag protein-tagged SIRT1, fulllength SIRT1 cDNA was amplified from a human cDNA library and fused at its C-terminus with sequences encoding monomeric protein. Briefly, the amplified SIRT1 fragment was inserted into the adenoviral vector, which contains the mouse cytomegalovirus (CMV) promoter, using the AdMax system. The resultant Flag-SIRT1 protein gene fusion was validated by nucleotide sequencing. Flag protein was detected by western blotting. Primary rat HSECs were transfected with this adenovirus vector to overexpress SIRT1, according to the manufacturer’s instructions. The SIRT1 sequences were used: sense (5-CGGGCCCTCTAGACTCGAGCGGCCGCATGATTGGCACCGATCCTC-3).

### Small interfering RNA (siRNA) transfection assay

Primary rat HSECs were transfected with siRNA to silence NOX2, p53 and progerin, according to the manufacturer instructions. The transfection efficiency was 75%. The following NOX2 siRNA sequences were used: sense (5-CCTCCTATGACTTGGAAAT-3). The following p53 siRNA sequences were used: sense (5-GGCTCCGACTATACCACTA-3). The following progerin siRNA sequences were used: sense (5-GCTCAGTGACTGTGGTTGA-3).

### Extraction of nuclear and cytoplasmic protein of primary rat HSECs

Nuclear and Cytoplasmic Protein Extraction Kit (Beyotime, P0028) was used to extract nuclear and cytoplasmic protein of primary rat HSECs (10^7^ cells per group). Nuclear and cytoplasmic protein was processed and detected for western blotting.

### Immunocytochemistry

Paraformaldehyde-fixed primary rat HSECs were incubated with primary antibodies, followed by the secondary antibodies, and subsequently mounted with DAPI. The primary antibodies included anti-NOX2 (1:200), anti-progerin (1:50), and anti-Ac p53 K381 (1:200). After incubation with primary antibodies and the secondary antibodies, HSECs were stained with the phallotoxin to detect F-actin. The number of positive cells was observed by fluorescence microscopy and quantified by Image J software.

### Co-immunoprecipitation (Co-IP)

Primary rat HSECs were transfected with SIRT1 adenovirus vectors, and were subsequently stimulated with H_2_O_2_ for 2 days. IP and immunoblotting (IB) were performed as previously described [5]. The antibodies for IP included anti-progerin and non-specific IgG; the antibodies for IB included anti-progerin, anti-Ac p53 K381, and anti-p53.

### Western blotting

Primary rat HSECs were isolated from normal rats and were treated with various stimulators, or were isolated from the model rats. HSECs were lysed in lysis buffer containing protease cocktails inhibitor and PMSF, as well as centrifuged at 12000r/min, 4°C, for 15min. The protein levels of HSECs were detected by western blotting. The primary antibodies included anti-vWF (1:1000), anti-NOX2 (1:1000), anti-NOX4 (1:1000), anti-Ac p53 K381 (1:1000), anti-p53 (1:1000), anti-progerin (1:25), anti-Lamin A/C (1:1000), anti-Lamin B1 (1:1000), anti-SIRT1 (1:1000), anti-Histone H3 (1:1000), and anti-GAPDH (1:1000). The secondary antibodies were HRP-conjugated Affinipure Goat Anti-Mouse IgG(H+L) (1:10000, Proteintech, SA00001-1) and HRP-conjugated Affinipure Goat Anti-Rabbit IgG(H+L) (1:10000, Proteintech, SA00001-2). The protein bands were visualized using the Pierce(tm) ECL Western Blotting Substrate.

## Statistical analysis

The data were reported as the mean ± standard deviation (SD) and were analyzed by SPSS17.0 software. In statistical analysis of two groups, a two-tailed Student’s t-test was utilized; whereas, in statistical analysis of more than two groups, one-way ANOVA was performed. P<0.05 was considered significant.

## Data availability

The datasets generated during and/or analyzed during the current study are available from the corresponding author upon request.

## Acknowledgements

The authors are much grateful to Mr. Litao Qin, Dr. Shasha Bian, and Dr. Yonghui Dong for essential helps in this study. This study is sponsored by WBE Liver Fibrosis Foundation (Grant no.CFHPC2019008), the National Natural Science Foundation of China (No. 81800551), the Scientific and Technological Project of Henan Province (No. 182102310210), and the Young Talent Research Cultivation Project of School of Clinical Medicine of Henan University (No.2019017).

## Author contributions

Xiaoying Luo designed the research, conceived ideas, performed experiments, wrote the manuscript, and obtained funding. Yangqiu Bai, Xiaoke Jiang, and Shuli He performed experiments and analyzed data. Zhiyu Yang, Di Lu, Suofeng Sun, Peiru Wei, Yuan Liang, Cong Peng, Ruli Sheng, Yaru Wang, Shuangyin Han, and Xiuling Li critically revised the manuscript. Bingyong Zhang designed the research, conceived ideas, and directed the study. All authors edited and reviewed the final manuscript.

